# The consumption of iceberg lettuce may reduce the adhesion of dietary fat to the mucus surface of the stomach barrier lining decreasing the risk of triggering acute gastroesophageal reflux

**DOI:** 10.1101/2023.08.26.554975

**Authors:** Thomas J. Hurr

**Affiliations:** South Australian Reflux Research Unit, Adelaide, South Australia, Australia

**Keywords:** lettuce, GERD, PPI, gastroesophageal reflux, contact angle, fat, trigger foods, lipophilic

## Abstract

Iceberg lettuce (IL) leaves contain surface epicuticular waxes that can adsorb significant quantities of dietary fat per unit weight. Fats such as olive oil, butter, lamb fat and lard readily adsorb onto IL with soft solid fats contained in lamb fat and lard, showing strong adhesion and more resistance to removal on rinsing in a water stream. Dishwashing detergents containing surfactants prevent the adsorption of both liquid and solid fats to the lettuce leaf surface, unless at very low concentrations, highlighting their significant influence on surface properties. Based on the adhesion of fats to the IL leaf surface, it is proposed the consumption of IL could reduce acute gastroesophageal reflux (GER) and the development of gastroesophageal reflux disease (GERD). In diets containing fat, the adhesion to IL reduces the amount of fat available to adhere to the mucus surface of the stomach barrier lining, protecting the lining and reducing access by trigger foods to underlying surfaces that could potentially initiate GER.

**Graphical abstract:** 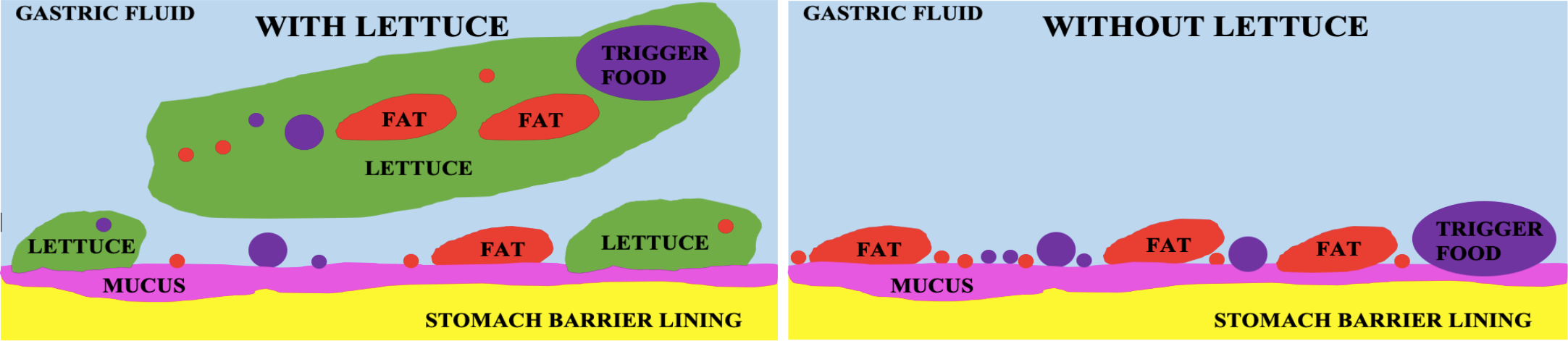

Iceberg lettuce leaves have approximately twice the surface area of the stomach for a 50g serve and 500cm^3^ meal volume and due to the waxy surface, can adsorb significant amounts of dietary fat. Lettuce may also adsorb on the mucus layer of the stomach barrier lining but is not regarded as a trigger food. Lettuce with its large surface area to adsorb fat, can keep fat suspended in solution, protecting the lining from fat adsorption and reducing the probability that fatty trigger foods can access underlying surfaces and cause acute gastroesophageal reflux.

## 1. Introduction

The diagnosis and management of gastroesophageal reflux (GER) and gastroesophageal reflux disease (GERD) for patients who have no alarm symptoms, commonly involves the use of proton pump inhibitors (PPIs) but doubts about their safety and long-term use have emerged [1]. Research into managing GERD through dietary choices rather than the use of PPIs is being undertaken, but it is not clear if any single key element in the diet is most essential [2]. A recent two case report in our research unit found consuming iceberg lettuce (IL) reduced the severity of acute GER but no explanation why this could occur, was given and so is undertaken in this report [3].

It is well established that eating leafy green vegetables is beneficial to health and it has been recommended 1 cup ≈ 75g, should be consumed daily [4]. It is also well established that diets high in fat, are both a potential risk in maintaining good health and in developing GERD [5-7]. Although total fat and vegetables have been considered individually, the fat/vegetable or fat/leafy green vegetable ratio was not found reported as an indicator to predict the risk factors of developing GERD. The dietary lipid/protein ratio and meal acidity has been considered as the most important factors associated with reflux and used to develop a scale for the refluxogenic potential of foods [8]. It is not well established why leafy green vegetables like IL should be consumed other than from the beneficial dietary components contained within, or the role surfactants (SF) may have in the vegetable/fat interactions present during digestion [9-13]. Although it is known that certain foods can trigger GER (trigger foods) and can have refluxogenic scores, the chemical and physical mechanisms of interaction with the mucus layer of the stomach barrier lining or underlying structures, that may initiate and maintain the reflux process, are unknown [8,14]. The relationship between dietary components and the effect on GERD symptoms is reportedly due primarily to decreased lower esophageal sphincter (LES) tone but includes transient LES relaxations, irritation of the esophageal mucosa, increase in gastric distention, altered gastric motility and increased gastric acid production [14]. Inflammation of the stomach barrier lining or underlying structures was not reported to be associated with acute GER or GERD [14].

A scanning electron microscopy study found leaf surfaces have complex epicuticular wax projections with various crystalloid structures that protect the leaf from environmental stresses [15]. The IL leaf surface has wax projections containing 70-85% 1-hexacosanol (C_26_ hydrocarbon chain length alcohol with melting point ≈ 76°C) distributed in varying amounts depending on both the location on a leaf and the leaf location within the lettuce [16]. The wettability of IL leaf surfaces is indicated by water contact angles (CA) from 60-100° showing both hydrophilic and hydrophobic character, where CA values between 0-90° indicate a hydrophilic surface and CA values greater than 90° indicate a hydrophobic surface [16,17].

Dietary fats, oils and waxes come from both plant and animal sources and include the glycerolipids as triacylglycerols, diacylglycerols and monoacylglycerols (fat), fatty acids, fatty alcohols, glycerophospholipids, sphingolipids, sterol lipids, prenol lipids, saccharolipids and polyketides [16,18]. The amount of dietary fat consumed each day is reported to be on average 81-87g/day with median values as saturated (32-35g/day), mono (28-31g/day) and polyunsaturated fats (12-13g/day) for both adult men and women with or without GERD [6]. Substituting monosaturated fat or polyunsaturated fat with saturated fats was not found to increase the risk of developing GERD [6]. Gastric lipases contribute to 10-30% of the overall adult dietary fat lipolysis and the stomach can absorb short and medium chain (<12 carbon atoms) fatty acids requiring 70-90% to be passed on to the small intestine for further digestion [19].

Surfactants are present in the digestive system to emulsify fats for transport and digestion [19-22]. Surfactants are found to occur naturally in foods, are added to commercially prepared foods, dishwashing detergents (DWD) and toothpastes [10-13]. Surfactants occur in saliva with a surface tension of 53-65mN/m, gastric fluid (43-54mN/m), the small intestine as bile salts (25-28mN/m which may also be present in gastric fluid) and in DWD (22-35 mN/m) [19-23]. An in vitro study to show how SF influence the digestibility of the fat tricaprylin, in the presence of the SF polysorbate 80 found that at low SF concentrations and in the presence of other types of SF (bile salts and phospholipids), adsorbed SF molecules can be displaced and allow lipolysis to occur in a concentration dependent manner [24]. Research involving the use of SF to prevent GERD was not found to be reported, possibly because lingual and gastric surfactants are not available for purchase and were not found included in standard (INFOGEST) in vitro gastric digestion methods [25]. This is not surprising given that the SF composition of saliva and gastric fluid are yet to be fully determined but are reported to consist of SF associated proteins A, B, C, D, fatty acids and triglycerides, pepsin, pulmonary SF like phospholipids involved in mucociliary clearance and possibly bile salts from the small intestine [19-22,26-28]. Investigation with DWD has found SF can remain on dishware after rinsing at concentrations of 0.15-0.4%, which can be above the concentration of SF required to emulsify fats in solution and can cause both gut epithelial inflammation, barrier damage and alter the gut microbiome [13,14]. As SF has a major impact on the surface properties of fat, are available in DWD and likely to be present in the human diet, DWD were used as the source of SF in the present study.

The gut barrier system is supported by complex protective mechanisms including mucus layers, a constantly renewing epithelial boundary, tight cell junctions and the gut microbiome [29]. The mucus layers are created by various *MUC* genes expressed throughout the body, which form mucin gels that coat all non-keratinized wet epithelial surfaces [29-33]. The stomach barrier lining consists of gastric epithelial cells coated with a dense mucus (mucins MUC5AC, MUC6) [32,33]. Particles with concentrated regions of positive charge or hydrophobicity, can bind and enrich in the outer mobile gastric mucus which is cleared rapidly once binding occurs [32]. It is also reported that particles in the trachea are formed into mucin MUC5B bundles which are then coated with mucin MUC5AC, as part of the mucociliary escalator and mostly swallowed [26]. Many visual models have been created to show how high fat diets increase intestinal permeability, modulate the composition and function of the mucus barrier layer, stimulating proinflammatory signalling, inducing epithelial cell stress and enriching the gut microflora [29-33]. Earlier visual models show the gastric mucus coated with a surface layer containing dipalmitoyl phosphatidylcholine (DPPC), an important pulmonary SF found in scrapings of the stomach wall and in the gastric fluid [27,28]. The DPPC surface layer was found to be hydrophobic with a water CA>90° in one model and with water CA 60-80° in another model, reduced after exposure to nonsteroidal anti-inflammatory drugs (NSAIDs) to a water CA<40° due to gastric inflammation [27,28]. If the gastric mucosal layer has hydrophobic surface regions due to the presence of pulmonary SF, then it is likely dietary lipophilic fat would be readily adsorbed (dispersive-dispersive attraction between hydrocarbon molecules) and possibly in a comparable way as found for the IL surface, which also has hydrophobic surface regions and a similar water CA≈ 60-100° [16,27,28,34].

This report investigates the relationship between IL/fat/SF adsorption and proposes that IL has a protective role by adsorbing dietary fat, reducing the probability that fat will adhere to the mucus surface of the stomach barrier lining, allowing mucosal integrity to be maintained for longer during the digestion of a fatty meal and lower the risk of developing GER and GERD.

## 2. Materials and methods

### 2.1. Foods and chemicals

The IL, olive oil (OO), butter (BU), lamb fat (LF) and lard (LD), water soluble food colouring, pasta (wheat lasagne sheets), apple (royal gala), cucumber (Lebanese), bread (wholemeal flour), minced lamb, cheese (mature), lean beef, spinach and 2 branded household DWD were purchased from local supermarkets. Lamb fat (LF) from different minced lamb samples, bought over 8 months, was extracted by heating to generate multiple samples of fat with several samples combined. Hydrochloric acid (HCl) and Sudan 111 (lysochrome dye) were purchased from Chem Supply (Adelaide). The IL was used directly as purchased without washing while the apples and cucumbers were washed first. Pasta was soaked in water for 30 minutes before use. The lean beef was pan fried without oil. Dish washing detergents as DWD1 contain coconut based anionic and non-ionic SF, DWD2 contain cationic (polyquaternium-7, cocamidopropyl betaine), anionic (sodium laureth sulphate) and non-ionic (capryloyl /caproyl methyl and lauroyl /myristoyl methyl glucamides) SF.

### 2.2. Hydrophilic/lipophilic dye solutions, rinsing and quantifying fat adsorption

All prepared solutions and rinsing or washing was carried out at 37°±3°C. To replicate a gastric solution, water or 0.1M HCl, without the inclusion of pepsin, mineral salts or lingual or gastric SF or lipases, was used [25]. To test the adhesive strength of the adsorbed fats to the IL leaf surface after agitation in prepared solutions for 30 seconds (s), the leaves were held vertically in air for 30s to determine how much fat (grams g) remains adhered under gravity and then rinsed in a water stream (WS) of running warm water for 30s at a flow rate of 42±2cm^3^/sec (1260±60cm^3^/30s) and again held vertically in air for 30s to again determine how much fat remains adhered. The amount of fat remaining adhered was termed the retention weight (RW). The WS was thought an aggressive way to determine the strength of IL/fat adhesion and although not the same process as gastric digestion, does allow visual observation of changes to the IL surface. The WS lacked careful temperature control, as solid LF and LD can have a softening or melting point around 37-38°C, with solid fat more resistant to removal and potentially increased the error in measuring the RW.

To prepare a solution to show the macroscopic hydrophilic nature of IL, approximately ≈ 0.5cm^3^ of green or blue food colouring dye was added to 90cm^3^ of 0.1M HCl. To prepare a solution to show the macroscopic lipophilic nature of the IL, approximately ≈ 0.005g of solid Sudan 111 (inclusion of solvents could dissolve epicuticular wax) was added to 10cm^3^ of either OO, BU, LF or LD to form a red liquid dye which can also contained some solids (LF or LD) and undissolved red dye particles. To determine the macroscopic hydrophilic/lipophilic surface properties of IL or other foods, the 2 dyed aqueous and fat solutions were mixed to form a solution containing 10% fat, shaken for 30s and allowed to stand for 10 minutes. At 37°C, the OO and BU were both liquids, LF consisted of both liquid and soft solids and LD consisted of both liquid and small particulate soft crystalline fat and with all fats, having a lower density than water, floating on the aqueous surface.

To determine the RW on the IL surface (after 30s under gravity and 30s in a WS), samples with the same surface area (SA) were required and cut from both inner and outer leaf regions of the IL as 5cm x 5cm = 25cm^2^ pieces. Samples with an initial weight (w_o_) were agitated for 30s in the solutions, removed, held vertically, allowed to drain in air for 30s and re-weighed (w_1_).

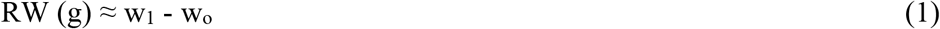

The samples were then held under a WS (standard deviation SD used for all numerical values of at least 3 repeat measurements) for 30s allowed to drain for 30s and re-weighed (w_2_).

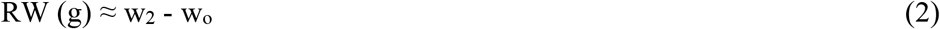

To calculate the average RW (ARW) so that comparison between samples could be made, the average initial weight w_o_(av) of IL samples with the same SA, was determined from 66 samples of 25cm^2^ pieces of IL leaf which varied from 0.99 g for the thinner outer leaf to 4.51g for the thicker inner leaf near the core with w_o_(av) ≈ 2.24±0.89g/25cm^2^.

To convert the RW of material adhered after dipping or rinsing, to ARW as g/100g IL, using the average weight of a 25cm^2^ section of IL;

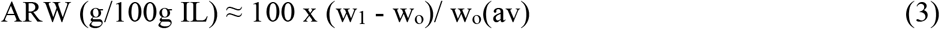

or for samples rinsed in a WS

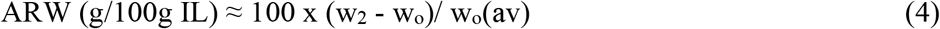

In this report, a region on the IL surface was arbitrarily assigned hydrophilic if it retained water, lipophilic if it retained fat and hydrophobic if it did not retain water, when removed from solutions containing water, fats, 0.1M HCl or DWD and allowed to drain in air for 30s [17,34]. Stable dry patches could also form on the IL leaf surface on the break-up of the solutions, which drained as rivulets, when the surfaces were held vertically, as noted previously for adiabatic surfaces and considered in this report as regions that do not show strong hydrophilic or lipophilic surface properties, under these conditions [35].

### 2.3. The use of dish washing detergents /surfactants

It was recognised that the surface properties of fats are greatly influenced by the presence of SF, even at very low concentrations and as human gastric SF is not available and at least some SF effects should be investigated, DWD was used. Commercial DWD contain SF at concentrations of ≈ 10-20% or 100-200g pure SF/litre so 0.1% DWD solutions used in this report, would contain 1g of the 10-20% DWD/litre or 0.1 – 0.2 g pure SF/litre [23]. Mixed SF solutions, as found in DWD, can have a critical micelle concentration (CMC) between 0.1-0.32g/litre [23]. The DWD that contain both amphoteric and non-ionic surfactant mixtures, can promote CMC lowering and the CMC values of the DWD was reported not to vary by more than 5% between 17-42°C [23].

To make the solutions containing DWD, pure fats were first dyed with ≈ 0.005g of solid Sudan 111 and then added to the 0.1M HCl/(0.1 – 0.5%) DWD solutions, shaken for 30s resulting in an emulsion of red fat droplets initially but within minutes formed a creaming layer (floating dispersed phase of the emulsion), allowed to stand for 10 minutes and to this solution the IL were added. If the fats were first shaken in water with 0.1-0.5% DWD1 or DWD2, the fat would not adsorb the solid Sudan 111 even after 8 hours, indicating the dye could not accessed lipophilic surface sites and that the fat was in a stable emulsion. For very dilute DWD solutions as 0.1 M HCl/10% OO/0.05% DWD, Sudan 111 would readily dissolve in OO despite the presence of DWD, to give a red floating coalesced layer of fat, indicating that a stable dispersion had not formed. No significant differences were observed when using either DWD1 or DWD2 when forming the fat solutions.

### 2.4. Hydrophilic /lipophilic nature of common food surfaces

To compare the adhesion of fat on IL with other common foods, intact samples IL, cheese, sliced apple, sliced cucumber, bread, pasta, dry pan fried beef and spinach samples (≈ 25cm^2^) were agitated in either 0.1M HCl/10% LF or 0.1M HCl/10% LF/0.1-0.5% DWD2 solutions for 30s, removed, held vertically and allowed to drain in air for 30s then agitated in water-bath (a WS would separate the bread matrix) for 30s to rinse any un-adsorbed fat from the surface, held vertically and allowed to drain in air for 30s again and re-weighed.

### 2.5. The retention weight of absorbed or adsorbed fats

To determine if water, acid solutions, SF or fats were absorbed into the interior of the intact IL leaf rather than adsorbed on the surface, samples were rinsed and then wiped with tissue paper and re-weighed to determine if an increase in the initial weight had occurred.

### 2.6. The ratio of lettuce:stomach surface area

The IL:stomach SA ratio should give an indication of the potential significance of IL as a competitive surface for the adsorption of fats. Magnetic resonance imaging of the stomachs of 12 healthy volunteers, after consuming 500cm^3^ of soup, found the maximum macroscopic stomach surface area (SA) = 536±25 cm^2^ (≈ 23 x 23cm) initially, decreasing over time, as the food was digested [36]. The macroscopic SA, using both sides of the IL leaf, can be calculated from w_o_(av) ≈ 2.24g/25cm^2^ such that the SA of 100g IL is (100/2.24) x 25 x 2 ≈ 2232cm^2^ (≈ 47 x 47cm) or for 1 serve of IL the SA is ≈ 1674cm^2^/75g IL (≈ 41 x 41cm) and for a smaller serve of 50g the SA ≈ 1116cm^2^/50g IL (33.5 x 33.5cm). The ratio of IL:stomach SA, for 1 serve of IL and a 500cm^3^ meal, is calculated as IL:stomach SA ≈ 1674:536 ≈ 3:1. For a smaller serve of 50g IL then IL:stomach SA = 1116:536 ≈ 2:1. If the meal size is reduced proportionally to the amount of IL consumed, the ratio of IL:stomach SA would be similar.

## 3. Results

### 3.1. Mapping the hydrophilic/lipophilic surface of intact IL

The intact IL leaf surfaces, when agitated in the dye solutions are shown to contain both hydrophilic (aqueous 0.1 M HCl dyed green), lipophilic (fat dyed red) and regions that appear smooth which do not hold either liquid (Figure 1). The aqueous hydrophilic green dye also absorbed into the cut surface edges of the IL, indicating a hydrophilic interior (Figure 1A-D). The use of mixed green and red dyed water/fat solutions to map surface hydrophilic/lipophilic surface sites was not found previously reported for food surfaces, only the use of water contact angles (sessile drop) to determine the local surface hydrophilic/hydrophobic nature. From the appearance of the IL leaf, the red dyed LF and LD were the more adhesive and more resistant to removal under a WS, compared to OO and BU (Figure 1).

**Figure 1:**
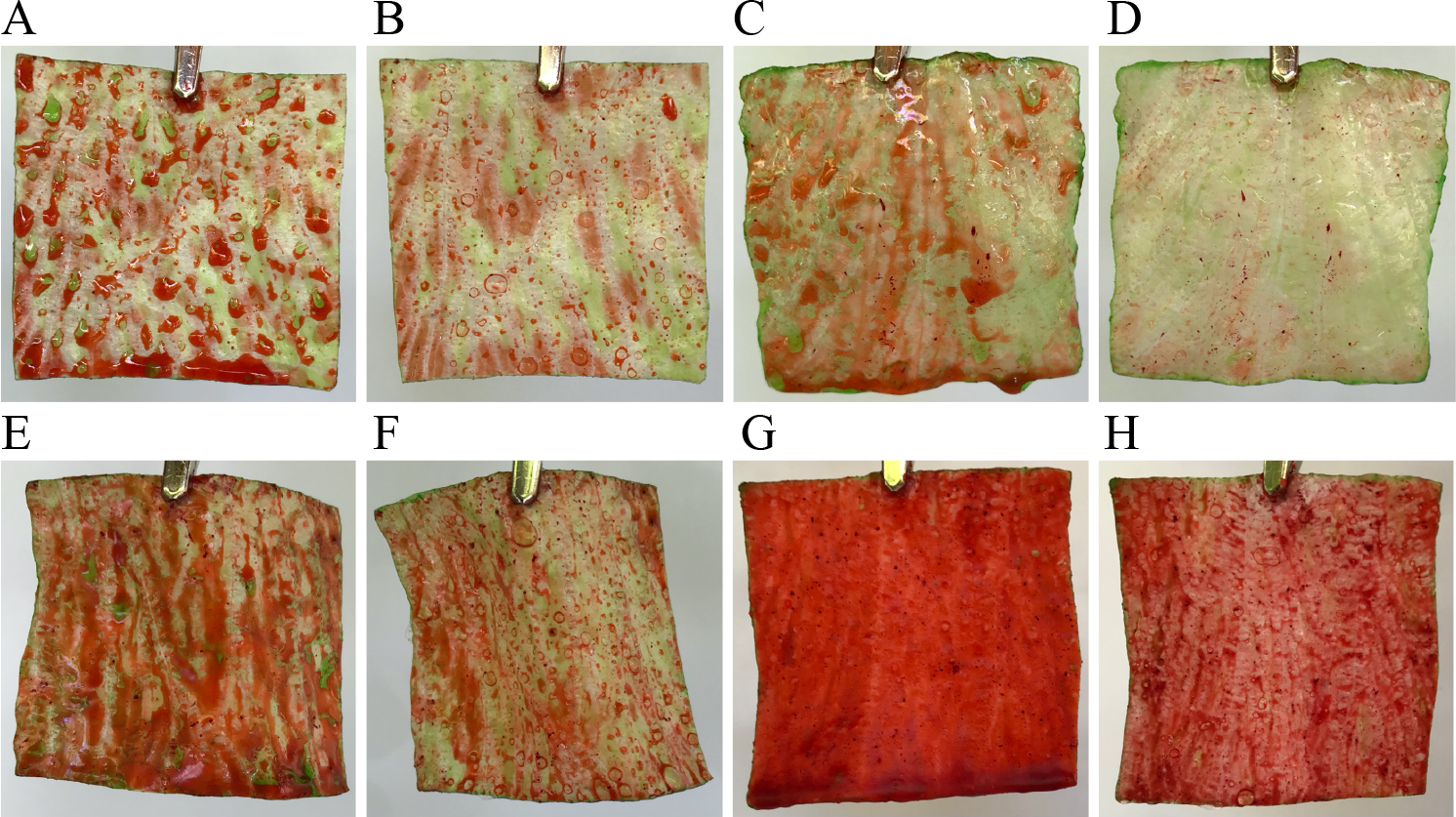
The adsorption of 0.1M HCl/10% fat solutions to the IL leaf surface showing hydrophilic (green) and lipophilic (red) surface sites after removal from solution and after rinsing in a WS. The weighed samples of IL (25cm^2^) were dipped in green dyed 0.1M HCl with 10% red dyed fat solution, agitated for 30s, then held vertically in air for 30s to drain, and re-weighed (**A, C, E, G**) followed by rinsing in a WS and again held vertically in air for 30s to drain and again re-weighed (**B, D, F, H**). **A**. OO, ARW = 17g/100g IL; **B**. OO, ARW = 6.7g/100g IL; **C**. BU, ARW = 20g/100gIL; **D**. BU, ARW = 5.8g /100g IL; **E**. LF, ARW = 28g /100g IL, **F**. LF, ARW = 10g /100g IL; **G**. LD, ARW = 41g/100g IL; **H**. LD, ARW = 8.9g/100 g. The ARW values show the combined weights of the adsorbed 0.1M HCl/fat solution but it can be seen from the amount of red dye, a significant proportion of the ARW value can be associated with fat adsorption particularly for the solid LF and LD.

### 3.2. The adsorption of fats to IL

The ARW values for the intact IL leaves are given for water, 0.1M HCl solution, pure fat (OO, BU, LF and LD) and 0.1M HCl/10% fat solutions includes the use of 0.05-0.5% DWD1 (Table 1). It is found that the ARW values for water (11±3.0g/100g IL) or 0.1M HCl (11±2.1g/100g IL) are similar, showing the adsorption of water to the IL surface was not significantly influenced by the ionic strength or pH of the solution and under the more dynamic conditions of a WS, the ARW are also similar, but lower (Table 1). The ARW values for pure liquid fats, OO and BU (ARW≈ 18-19±4.0g/100g IL) were also surprisingly similar for OO and BU in 0.1M HCl (ARW≈ 17-18±2.9g/100g IL) without the expected increase in weight due to the additional adsorption of 0.1M HCl (Figure 1A, 1C, Table 1). This result indicates the 0.1M HCl solution may change the surface properties of OO, BU or IL possibly by restructuring the behaviour of specific surface molecules selectively exposing hydrophilic or lipophilic parts [34].

**Table 1:**
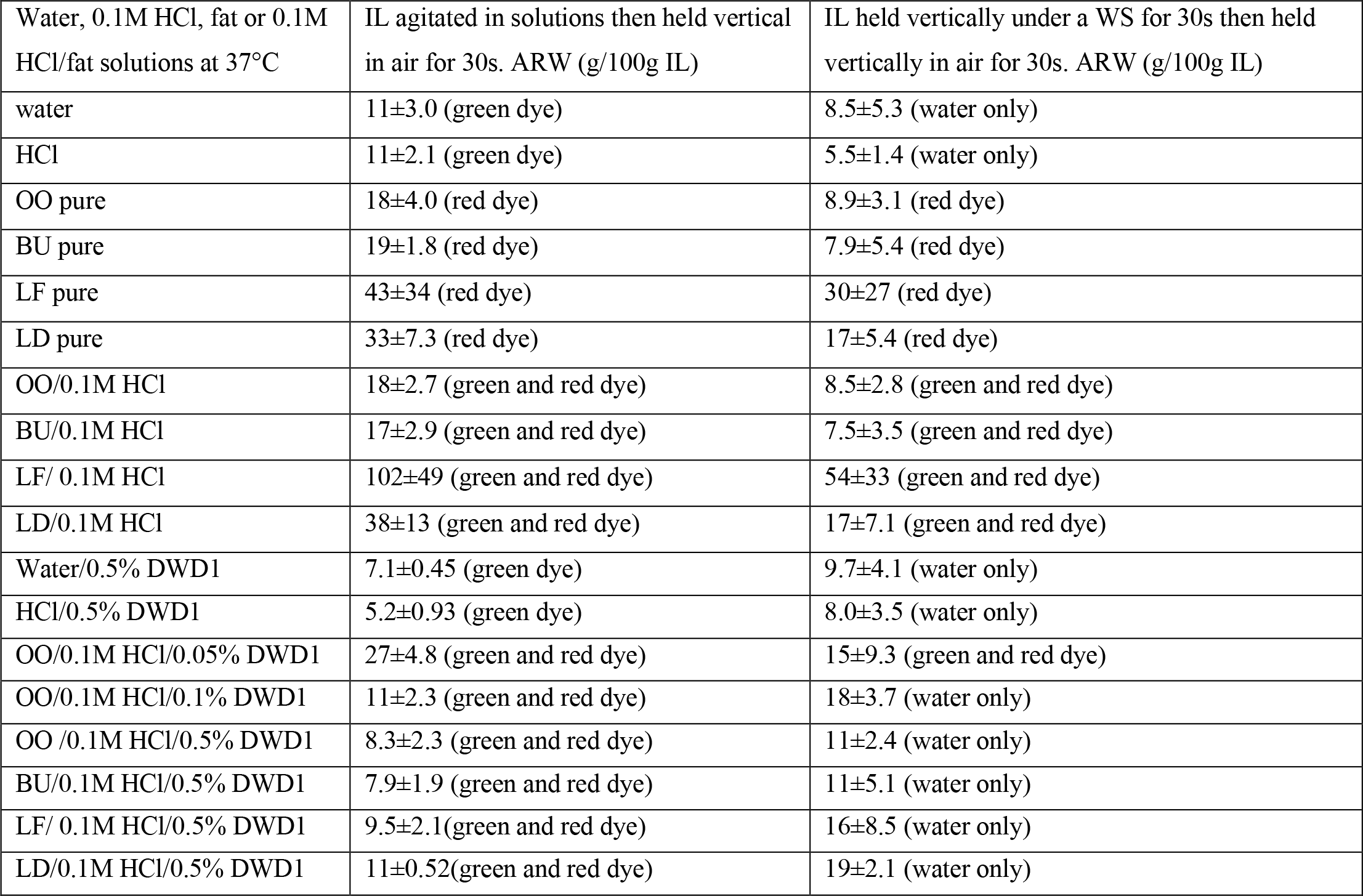
The strength of water and fat adhesion to IL is indicated by ARW (g/100g IL) for samples held vertically in air and under a WS (equations 3 and 4). The liquids used were water, 0.1M HCl, pure fats (OO, BU, LF and LD), fats as 10% solutions with 0.1M HCl with or without DWD1. The IL was agitated in the solutions for 30s, removed, held vertically in air for 30s and allowed to drain and re-weighed then held under a WS for 30s, then again allowed to drain for 30s and again re-weighed. The ARW values for the pure fats (without water) indicate fat only retention prior to rinsing in a WS, all other ARW values include the weight of adsorbed water or DWD1, with the amount of dyed fat adsorbed being visually apparent. After rinsing in a WS, green dye is removed but dyed fats (red) can remain adsorbed, with the fats containing solids (LF and LD) showing the greatest resistance to removal (supplementary material).

| Water, 0.1M HCl, fat or 0.1M HCl/fat solutions at 37°C | IL agitated in solutions then held vertical in air for 30s. ARW (g/100g IL) | IL held vertically under a WS for 30s then held vertically in air for 30s. ARW (g/100g IL) |
| --- | --- | --- |
| water | 11±3.0 (green dye) | 8.5±5.3 (water only) |
| HCl | 11±2.1 (green dye) | 5.5±1.4 (water only) |
| OO pure | 18±4.0 (red dye) | 8.9±3.1 (red dye) |
| BU pure | 19±1.8 (red dye) | 7.9±5.4 (red dye) |
| LF pure | 43±34 (red dye) | 30±27 (red dye) |
| LD pure | 33±7.3 (red dye) | 17±5.4 (red dye) |
| OO/0.1M HCl | 18±2.7 (green and red dye) | 8.5±2.8 (green and red dye) |
| BU/0.1M HCl | 17±2.9 (green and red dye) | 7.5±3.5 (green and red dye) |
| LF/ 0.1M HCl | 102±49 (green and red dye) | 54±33 (green and red dye) |
| LD/0.1M HCl | 38±13 (green and red dye) | 17±7.1 (green and red dye) |
| Water/0.5% DWD1 | 7.1±0.45 (green dye) | 9.7±4.1 (water only) |
| HCl/0.5% DWD1 | 5.2±0.93 (green dye) | 8.0±3.5 (water only) |
| OO/0.1M HCl/0.05% DWD1 | 27±4.8 (green and red dye) | 15±9.3 (green and red dye) |
| OO/0.1M HCl/0.1% DWD1 | 11±2.3 (green and red dye) | 18±3.7 (water only) |
| OO /0.1M HCl/0.5% DWD1 | 8.3±2.3 (green and red dye) | 11±2.4 (water only) |
| BU/0.1M HCl/0.5% DWD1 | 7.9±1.9 (green and red dye) | 11±5.1 (water only) |
| LF/ 0.1M HCl/0.5% DWD1 | 9.5±2.1 (green and red dye) | 16±8.5 (water only) |
| LD/0.1M HCl/0.5% DWD1 | 11±0.52 (green and red dye) | 19±2.1 (water only) |

For pure LF, the ARW ≈ 43±34g/100g IL has a large standard deviation, reflecting variations in the amounts of fat that adsorbed from mixtures of both liquid and solid LF, with lower values resulting from predominantly liquid fat adsorption, which could drain from the surface when held vertically, to higher values when solid fats adsorb, that resisted draining. Solid LF was also more readily retained on rinsing in a WS (ARW≈ 30±27g/100g IL) than the liquid only fats (OO and BU, Table 1). Lamb fat from the LF/0.1M HCl solutions could adsorb as a large soft solid ball (ARW ≈ 102±49g/100g IL) which resists removal in a WS (ARW ≈ 54±33g/100g IL, Table 1). The variations in the amount of LF that adsorb, may be due to the range in the melting points for animal fat, which depend on the origin from the animal body, with back fat 30-40°C, leaf fat 43-48°C and mixed fats 36-45°C [37]. Lard, like LF, is composed of both liquid and solid fats with ARW≈ 38±13g/100g IL and after rinsing in a WS, ARW≈ 17±7.1g/100g IL for the LD/0.1M HCl solutions but with lower standard deviation values, due to the more consistent composition of a commercial product than for LF, extracted from different mince samples.

The ARW values for IL agitated in water/0.5% DWD1 (7.1±0.45g/100g IL) or in 0.1M HCl/0.5% DWD1 (5.2±0.93g/100g IL) were similar but lower than the ARW when only water or 0.1M HCl solutions were present (Table 1). The lower ARW values when DWD1 was present may be due to the difference in surface tension between water and water/SF solutions, with the lower surface tension SF solutions draining from the IL surface more efficiently. When water/0.5% DWD1 and 0.1M HCl/0.5% DWD1 were rinsed in a WS, the ARW values increased, possibly due to increased wetting of the surface regions between the epicuticular waxes.

The ARW values for IL agitated in 0.1M HCl/10% fat (OO, BU, LF, LD)/0.1-0.5% DWD1 solutions and then held vertically in air, were lower than those in the absence of DWD1, with fat adsorbing as part of an emulsion and readily rinsed from the IL surface within seconds in a WS. The same results were also found for 0.1M HCl/10%OO/0.1% DWD2 solutions, where on rinsing in a WS, all adsorbed dyed OO was removed from the IL leaf surface (Figure 2A, 2B). Only when the DWD1 was at a very low concentration of 0.05% could dyed OO adsorb on the IL leaf surface and remain adsorbed, after rinsing in a WS, indicating the significant influence of SF on IL/fat interactions (Table 1). Epicuticular wax structures remained intact on the IL despite rinsing in DWD2 as OO could re-adsorb from a 0.1M HCl/10%OO solution on to the IL surface (Figure 2C-2G). Scanning electron microscopy (SEM) images of soybean leaves show SF solutions can partially remove some but not all epicuticular wax from these leaf surfaces [38].

**Figure 2:**
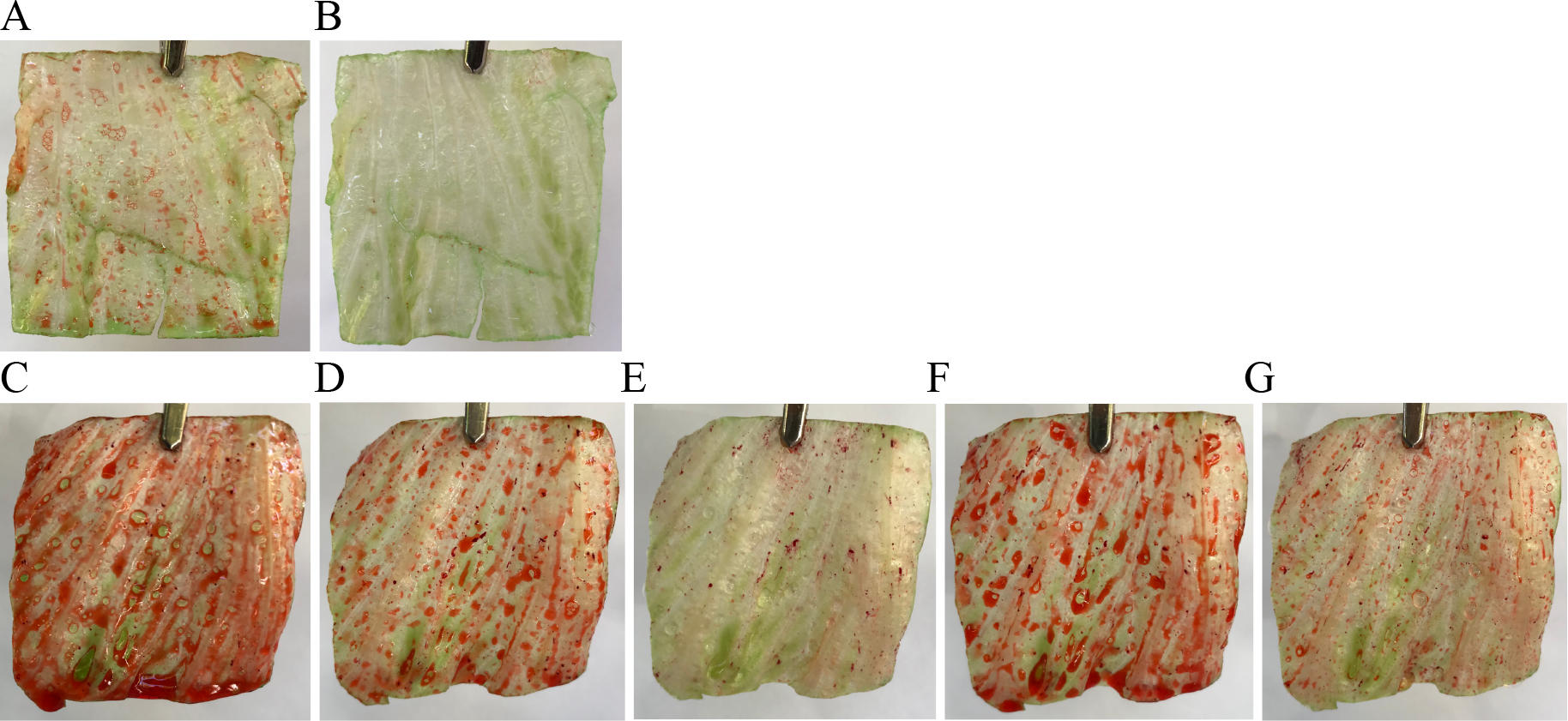
The influence of OO adsorption to IL from OO/DWD2 solutions (A-B) and on rinsed in DWD2 (C-G). **A**. a sample of IL (≈ 25 cm^2^) agitated for 30s in green dyed 0.1M HCl/ red dyed 10%OO/0.1% DWD2, then held vertically in air for 30s, **B**. the sample in (A) was then rinsed in a WS and allowed to drain for 30s. The sequence A-B shows OO weakly adsorbed from DWD2 solution and is easily rinsed from the IL surface. **C**. a sample of IL (≈ 25 cm^2^) agitated for 30s in green dyed 0.1M HCl/ red dyed 10%OO (no DWD2), then held vertically in air for 30s showing both water and OO adsorption, **D**. the sample shown in (C) was agitated in DWD2 for 30s held in air and allowed to drain for 30s showing some OO was removed. **E**. the sample shown in (D) was then rinsed in a WS held in air and allowed to drain for 30s showing the OO rinsed away with red particles on the IL surface from undissolved solid Sudan 111. **F**. the sample shown in (E) was re-agitated in the original 0.1M HCL/10%OO solution (no DWD2) used to generate sample C, held in air and allowed to drain for 30s shows OO can re-adsorb. **G**. the sample in (F) was re-rinsed in a WS then held in air and allowed to drain for 30s showing most of the OO rinsed away. The sequence (C-G) shows that OO adsorbed to the IL surface in (C) is partly removed from the surface by DWD2 in (D) and completely removed in a WS in (E) but DWD2 does not remove the surfaces epicuticular waxes as shown in (F) as OO can re-adsorbs on to the IL surface with residual OO still remaining after rinsing in a WS in (G).

### 3.3. Fat adsorption or absorption

The water, pure fats, fats/0.1M HCl and fats in DWD solutions that had adsorbed onto the IL and were liquids at 37°C, could be wiped from the IL leaf surface using tissue paper (ARW≈ 0g/100g IL) showing water, fats and DWD did not significantly absorb into the bulk of the IL.

### 3.4. The adsorption of fats on common food surfaces

The IL, cheese, apple, cucumber, bread, pasta, meat and spinach were agitated for 30s in a blue dye solution of 0.1M HCl/10% LF dyed with Sudan 111, held vertically in air for 30s, allowed to drain and washed in a water bath at 37°C (Figure 3). A LF sample that strongly adhered to IL was chosen to test the adhesion of fat to the other food surfaces (Figure 3A). The macro-nutrient composition and assigned hydrophilic or lipophilic surface properties of the foods, based on visual observations of the amounts of dye adsorbed, are shown in Table 2 [39]. The IL leaf strongly adsorbed LF and this was not easily washed from the surface in a water-bath (ARW ≈ 53g/100g IL, Figure 3A). Cheese adsorbed mainly aqueous blue dye indicating a mostly hydrophilic surface with some small lipophilic red patches (Figure 3B). The lipophilic apple and cucumber skins could adsorb LF, but the fat could readily be wiped from the smooth skin surface (Figure 3C, 3D). Bread showed a strong adsorption of blue dye in the inner regions and red dyed LF on the crust where fat (grease) had been used on the baking container, while pasta had a completely hydrophilic surface (Figure 3E, 3F).

**Table 2:** Values for the macro-nutrient composition of foods from Food Standards Australia and New Zealand, Australian food composition database [39] and the classification of food surface properties based on the appearance of dyes after adsorption from solutions containing both blue dyed 0.1M HCl, showing hydrophilic surface regions and red dyed LF, showing lipophilic surface regions (Figure 3). The weight % values may not add to 100% due to minerals and other micro-components. ^1^Note IL does not have a value recorded for fat as amounts are presumably not significant for macro-nutrient compositions. ^2^g/100cm^3^ converted to g/100g; ^3^estimated values.

| Food | Water weight % | Carbohydrate (starch, sugars and fibre) weight % | Protein weight % | Fat weight % | Classification of surface properties using dyed aqueous /fat solutions |
| --- | --- | --- | --- | --- | --- |
| iceberg lettuce | 96 | 2 | 0 | 0<sup>1</sup> | hydrophilic /lipophilic |
| cucumber (Lebanese) | 96 | 3 | 0 | 0 | hydrophilic except skin |
| spinach (fresh raw) | 92 | 2.5 | 3 | 0 | hydrophilic /hydrophobic |
| apple (royal gala) | 87 | 12 | 0 | 0 | hydrophilic except skin |
| pasta (white wheat) | 63 | 28 | 5 | 0 | hydrophilic |
| bread (wholemeal flour) | 39 | 46 | 10 | 2-3 | hydrophilic |
| cheese (mature) | 34 | 0 | 25 | 34 | hydrophilic /lipophilic patches |
| beef (dry fried) | 65 | 0 | 32 | 3 | hydrophilic |
| olive oil (OO) | 0 | 0 | 0 | 100<sup>2</sup> | lipophilic |
| butter (BU) | 15 | 0 | 1 | 81 | lipophilic |
| lamb fat (LF) | 0 | 0 | 0 | 100<sup>3</sup> | lipophilic |
| lard (LD) | 0 | 0 | 0 | 100<sup>3</sup> | lipophilic |

**Figure 3:**
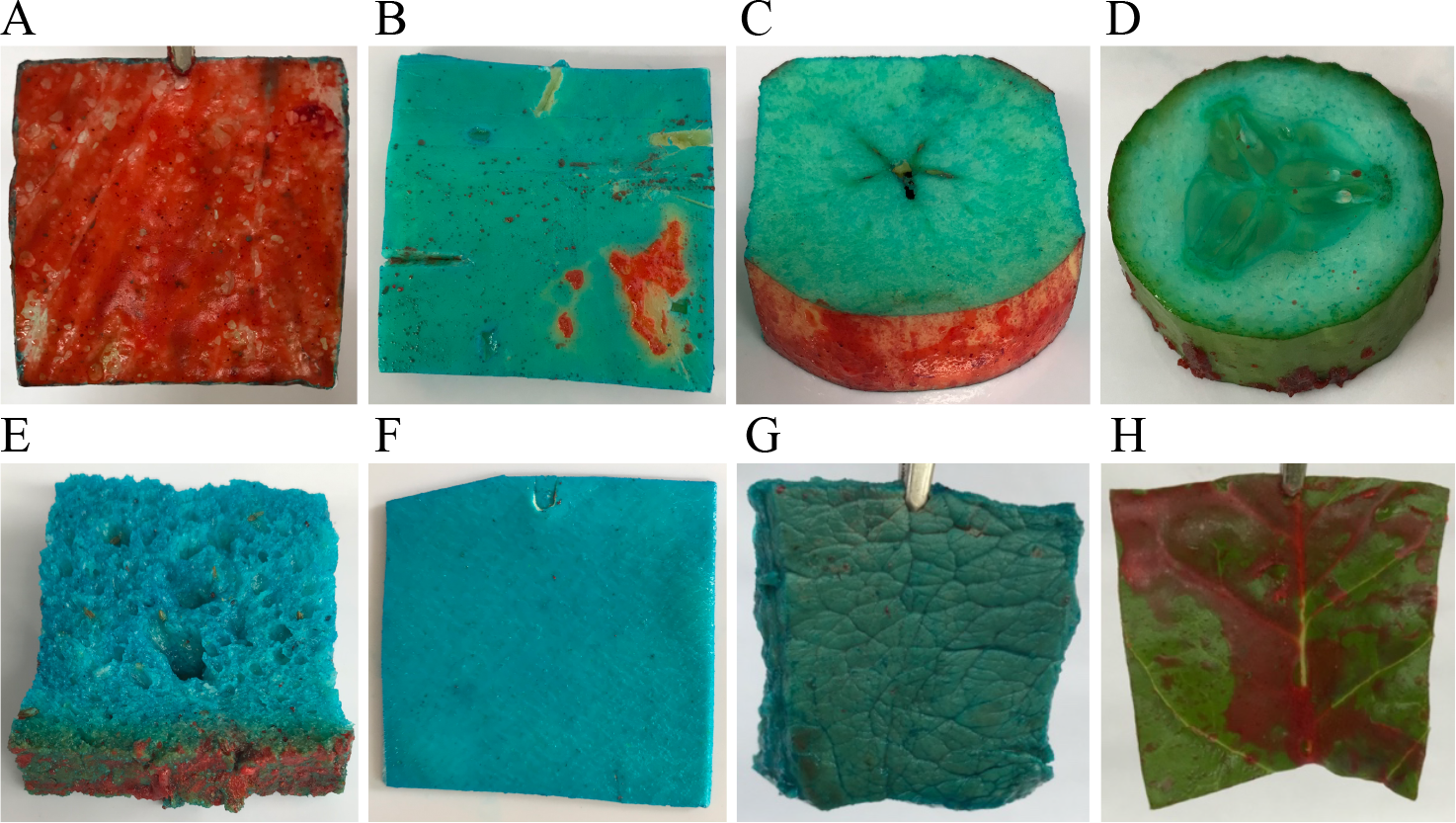
Common foods (≈ 25cm^2^) agitated in aqueous 0.1M HCl(blue)/10% LF (red) solution for 30s, allowed to drain in air for 30 s, washed in a water bath for 30s, then held in air and allowed to drain for 30s, at 37°C. The adsorption of aqueous blue dyed 0.1M HCl solution shows hydrophilic surface regions and red dyed LF show the lipophilic surface regions. **A**. IL, ARW ≈ 53g/100g IL after washing, **B**. cheese, **C**. apple, **D**. cucumber, **E**. bread, **F**. pasta, **G**. fried beef and **H**. spinach, RW ≈ 59g/100g spinach after washing (RW = w_2_-w_o_/w_o_ where w_o_ = 1.29g, w_2_ = 2.05g).

The dry pan-fried beef was found to be hydrophilic despite containing protein, known to consist of amino acids, some of which contain hydrophobic chemical groups [40]. Spinach, like IL, has a surface which contains epicuticular waxes and was found to adsorb LF in patches with some smooth sections that did not strongly adsorb either water or fat (Figure 3H). If DWD2 (0.1-0.5%) was included in the fat solutions, none of the foods adsorbed significant amounts of LF after washing in a water bath.

## 4. Discussion

This study has demonstrated that by using mixed dyed aqueous/fat solutions, the hydrophilic/lipophilic surface properties of food can be identified (Figure 1-3). Raw leafy green vegetables like lettuce with an outer protective layer of epicuticular waxes, can adsorb significant quantities of dietary fat. Apart from spinach, other common foods tested do not maintain the adsorption of any significant amounts of dietary fats from the 0.1M HCl /fat solutions after rinsing in a water bath, despite some containing protein and fat (cheese and beef, Table 2, Figure 3).

Intact IL can adsorb dietary fats, with soft solid LF and LD showing higher ARW values and greater resistance to removal under a WS, than the liquid fats, OO or BU at 37 °C. (Figure 1, Table 1). The ARW values for the 4 pure fats (OO, BU, LF, LD, drained in air for 30s) can be added together as 18+19+43+33 ≈ 28.2g/100g IL or 21g/75gIL to give an average value for lipophilic fat adsorption to IL (Table 1). If the average consumption of fat is ≈ 81-87g/day then 1 serve of IL (75g) could adsorb ≈ 21/84 ≈ 25% of average daily fat consumption [6]. Although only a very approximate result and the influence of dietary SF unclear, it still indicates the potential significant value IL may have in adsorbing and suspending lipophilic dietary fat as fat/IL particles in the gastric fluid, reducing fat exposure and possible adhesion or absorption to the mucus surface of the stomach barrier lining. In the presence of DWD, both liquid and solid fats poorly adsorb on to the IL leaf surface and the fats that did adsorb, were readily removed on rinsing under a WS unless the concentration of DWD was very low (Table 1, Figure 2A, 2B). It was unexpected that the SF in DWD, designed to clean surfaces, would be so powerful in preventing the adsorption of both liquid and solid fats to the IL leaf surface, even at low concentrations, suggesting studies into the adsorption of fats to food surfaces should consider the influence of SF (Figure 2A, 2B).

A model, showing how fat and trigger foods interact in the presence and absence of IL with the mucus surface of the stomach barrier lining, is proposed (graphical abstract). If 50g of IL in the diet is consumed with a meal of volume 500cm^3^, then the IL has approximately twice the SA as the stomach and so can potentially adsorb twice as much dietary fat, assuming the kinetics of adsorption are similar, which may be probable given the gastric mucosal surface may contain an outer hydrophobic layer with a similar CA as the IL surface [16,27,28]. The waxy surface of IL may adsorb on the mucus surface of the stomach barrier lining, but IL is not recognised as a trigger food and due to the large lipophilic surface area of IL, reduce the amount of fat that can adsorb on the mucus layer. The IL/fat and fat that adhere and slough off the mucus surface together with the IL/fat particles that do not adhere to the mucosal surface, progress to the small intestine to be digested by bile salts and lipases [22,32]. Benefits of consuming IL may also extend to diets that consume foods with low natural SF content or lack sufficient lingual or gastric SF to emulsify fats, which could otherwise reduce the concentration of adhesive fat deposits on the gastric mucosal surface. If commercially prepared SF from DWD were present, the fats would be dispersed in the gastric solution, but SF may also damage the mucosal layer and potentially the underlying epithelial cells together with changes to the mucosal microbiome [12,13]. It is likely SF have an important role in the digestion of fat in the gastric environment but must be of the appropriate chemical composition and concentration to be beneficial [31,41].

It is still unknown if the interactions between lettuce, fat, SF and the mucus surface have a role in reducing GER or GERD. Further studies could consider particulate IL leaves, fat/SF/gastric mucosa interactions, gastric SF, the gut/lung axis and the role of lipases.

## 5. Conclusion

The large lipophilic surface area per unit weight of IL, lacking in other common foods, gives IL the potential to adsorb significant quantities of dietary fat, with solid fats more resistant to removal on rinsing in water than liquid fats. It was found that DWD can prevent the adsorption of fat to the IL leaf surface, except at very low concentrations, indicating a potentially significant role of SF in dietary fat/food interactions. An explanation why IL in the diet could reduce GER and GERD is proposed, based on the competitive adsorption of dietary fats between the IL leaf and the stomach barrier lining. The strong adhesion of solid fats to IL may allow fats to pass from the gastric environment to the small intestine, minimizing irritation and adhesion to the mucus layer of the stomach barrier lining, lowering the probability trigger foods can access underlying structures and cause GER and GERD. The ability of IL surfaces to adsorb dietary fats may prove to be a supporting factor in emerging evidence that the consumption of vegetables, which includes lettuce and leafy vegetables, can be negatively correlated with the risk of developing GERD.

## Abbreviations

IL: iceberg lettuce
SF: surfactant
CA: contact angle
OO: olive oil
BU: butter
LF: lamb fat
LD: lard
DWD: dishwashing detergent
s: second
WS: water stream
RW: retention weight
ARW: average retention weight
SA: surface area
GER: gastroesophageal reflux.

## Supplementary Data

Supplementary data is available on request.

